# Inferring secretory and metabolic pathway activity from omic data with secCellFie

**DOI:** 10.1101/2023.05.04.539316

**Authors:** Helen O. Masson, Mojtaba Samoudi, Caressa M. Robinson, Chih-Chung Kuo, Linus Weiss, Km Shams Ud Doha, Alex Campos, Vijay Tejwani, Hussain Dahodwala, Patrice Menard, Bjorn G. Voldborg, Susan T. Sharfstein, Nathan E. Lewis

## Abstract

Understanding protein secretion has considerable importance in the biotechnology industry and important implications in a broad range of normal and pathological conditions including development, immunology, and tissue function. While great progress has been made in studying individual proteins in the secretory pathway, measuring and quantifying mechanistic changes in the pathway’s activity remains challenging due to the complexity of the biomolecular systems involved. Systems biology has begun to address this issue with the development of algorithmic tools for analyzing biological pathways; however most of these tools remain accessible only to experts in systems biology with extensive computational experience. Here, we expand upon the user-friendly CellFie tool which quantifies metabolic activity from omic data to include secretory pathway functions, allowing any scientist to infer protein secretion capabilities from omic data. We demonstrate how the secretory expansion of CellFie (secCellFie) can be used to predict metabolic and secretory functions across diverse immune cells, hepatokine secretion in a cell model of NAFLD, and antibody production in Chinese Hamster Ovary cells.

## Introduction

Protein secretion is fundamental to all living organisms. The mammalian secretory pathway processes approximately one third of all protein coding genes (Gutierrez et al., 2020; Uhlén et al., 2015) including both secreted and membrane-bound proteins responsible for controlling cell-cell communication, regulating cell adhesion, and modifying the extracellular environment. Unsurprisingly, perturbations to the secretory system are seen in many diseases including cancer (Glavey et al., 2015), Alzheimer’s (Bharadwaj et al., 2010), and Parkinson’s (Winklhofer and Tatzelt,). Protein secretion is also heavily exploited in biotechnology to manufacture biopharmaceuticals and industrial enzymes. Due to its widespread importance, understanding protein secretion is fundamental to elucidating the underlying pathophysiology of diseases, and is paramount for the efficient production of life-altering biotherapeutics.

While advances in high-throughput technologies and sequencing methods have enabled comprehensive monitoring of biological systems with omics, complex interdependencies between molecular components and inadequate mechanistic understanding of the system limits our capacity to interpret omic data. To this end, numerous approaches strive to identify mechanisms linking gene and/or protein expression patterns to phenotype using known biological pathways and feature engineering. Many of these methods aim to reduce complexity of the system by grouping genes into “functional clusters” representative of a particular pathway or biological process, often followed by statistical enrichment analyses (Huang et al., 2009; Khatri et al., 2012). Although these platforms accurately capture changes in biological pathways, their ability to capture cell functions is limited as they often lack the mechanistic information underlying these changes in phenotype. Accordingly, researchers have developed genome-scale models as tools to interpret large omic data sets (Hyduke et al., 2013). These models provide a mechanistic link between gene expression and phenotype, however they require extensive computational knowledge and experience in systems biology to deploy.

To increase accessibility of complex systems biology analyses enabled by genome-scale models a computational toolbox called CellFie (**Cell F**unction **i**nferenc**e**) was developed (Richelle et al., 2021). The CellFie toolbox precomputes metabolic pathway activities involving thousands of reactions (Hefzi et al., 2016; Sigurdsson et al., 2010; Swainston et al., 2016) to identify biologically interpretable gene modules (“tasks”) within the genome-scale network (Richelle et al., 2019a). Complementing current enrichment analyses, the algorithm overlays omic data onto the precomputed modules to quantify the consistency of protein/gene expression with the ability to accomplish each task (Richelle et al., 2019b).

Here, we broaden the scope of CellFie to include cell functions involved in protein secretion, which we call secCellFie. Thanks to the recent development of genome-scale models of mammalian protein secretion (Gutierrez et al., 2020), we curated a set of 21 tasks covering core secretory pathway functions. These functions are integrated with the CellFie algorithm to enable the prediction of not only metabolic cell function, but also protein secretion capabilities. We demonstrate the broad utility of this tool with a series of vignettes. First, we demonstrate how the addition of secretory tasks enhance CellFie’s ability to cluster immune cell types using gene expression data from the Human Protein Atlas (HPA) (Uhlén et al., 2015). Second, we show an industrial application of secCellFie comparing high and low monoclonal antibody (mAb)-producing CHO cells. Finally, we apply our method to RNA-Seq data in the study fetuin-B secretion, a key hepatokine involved in non-alcoholic fatty liver disease (NAFLD), and demonstrate more broadly how the altered processes are connected by using *in situ* protein-protein interaction data, as measured using the Biotinylation by Antibody Recognition (BAR) (Bar et al., 2018). These case-studies highlight how analyzing omic data with model-driven functional modules of the secretory pathway can enhance biological insight and drive hypothesis generation.

## Materials and Methods

### Curating secretory tasks

CellFie tasks consist of groups of reactions that work together to complete a biological function. The secretory tasks presented here were manually curated from the genome-scale reconstructions of mammalian secretory pathways (Gutierrez et al., 2020) based on extensive literature and database research. Manual curation resulted in a collection of 21 secretory tasks associated with 7 systems (translocation, processing in the ER, proteostasis, processing in the Golgi, and vesicle trafficking). A full table of secretory tasks and their corresponding reactions for human, mouse, and CHO can be found in Supplementary Data 1.

### Integrating genome-scale models with secretory pathway models

CellFie relies on the mechanistic links provided in genome-scale models to infer task activity from omic data. We utilized the data and code from Gutierrez et al. (Gutierrez et al., 2020) to integrate the secretory reactions of CHO, mouse, and human into genome-scale models iCHOv1 (Hefzi et al., 2016), iMM1415 (Sigurdsson et al., 2010), and Recon2.2 (Swainston et al., 2016), respectively. The resulting models combining metabolism and protein secretion can be found on <u>GitHub</u>.

### Quantifying secretory tasks from omic data

We used the previously described task framework (Richelle et al., 2021) to calculate activity scores for the 21 secretory tasks. We provide users the option to run CellFie for metabolic, secretory, or combine metabolic and secretory analyses.

### HPA immune cell CellFie analysis

Protein-coding transcripts per million (pTPM) data from HPA was preprocessed into an expression matrix compatible with CellFie: (i) ensemble gene IDs were converted to entrez IDs using the AnnotationDbi package (Pagès et al., 2020) in R, (ii) duplicate gene entries were aggregated based on max gene expression, and (ii) a pseudo count of 1e-05 was added. We used this preprocessed expression matrix to quantify metabolic and secretory scores using the secretory expansion of genome-scale model Recon2.2 (Swainston et al., 2016) with the following parameters: local minmaxmean threshold with upper and lower percentiles of 25 and 75 respectively. Hierarchical clustering of tasks scores was performed and visualized using the pheatmap package (Kolde, 2019) in R. Dendrograms were compared using FM-Index correlation from the dendextend package (Galili, 2015) and visualized with the corrplot package (Wei and Simko, 2021) in R. Hypergeometric enrichment of CellFie pathways in each DC cluster was carried out using the clusterProfiler package (Yu et al., 2012) in R. Lastly, differentially active tasks between DC subtypes were calculated using Welch’s t-test. Significant tasks (|FC| ≥ 1.25, FDR < 0.1) were visualized in a volcano plot using the EnhancedVolcano package (Blighe et al., 2021) in R.

### Cell culture and generating the IgG producing CHO cells

Recombinant monoclonal humanized IgG CHO cell lines were developed and cultured as previously reported (Dahodwala et al., 2019; Jiang et al., 2006; Jiang and Sharfstein, 2009). Briefly, CHO cells were co-transfected with two plasmids, one containing IgG heavy chain and DHRF genes and the other containing IgG light chain and neomycin phosphotransferase (Neo) genes. Transfected cell lines were initially selected using 400 µg/mL Neo, and all subsequent cultures were performed in the absence of Neo. Gene amplification was then performed by stepwise selection with increasing concentrations of MTX. For this study, a lower producing parental cell line A0 and its DHFR/MTX-amplified high-producing progeny A1 were chosen for investigation.

### CellFie analysis of IgG producing CHO cells

Expression data was run through secCellFie using the secretory expansion of genome-scale model MT_iCHOv1_final (Hefzi et al., 2016) with the following parameters: local minmaxmean threshold with upper and lower percentiles of 25 and 75 respectively. Samples CHO-Core-8 (A0_EXP) and CHO-Core-16 (A1_STA) appeared to be skewed (Figure S2), thus these samples were normalized by dividing CellFie scores by the slope of the best fit line (Figure S3). Principal component analysis was performed in R and visualized using the factoextra package (Anon, 2020). Differential expression analysis to identify differentially expressed GPI-anchor proteins was performed using DESeq2 package (Love et al., 2014) in R. Genes encoding GPI-anchor proteins were identified based on their annotation in the Protein Specific Information Matrices (PSIM) (Gutierrez et al., 2020). Significantly differentially expressed (|FC| ≥ 1.5, FDR < 0.1) GPI-anchor proteins were visualized using the gplots package (Warnes et al., 2022) in R. Differentially active tasks were determined using Welch’s t-test. Select secretory (FDR < 0.1) and metabolic (|FC| ≥ 1.5, FDR < 0.1) tasks were visualized using the ggplot2 package (Wickham, 2016) in R.

### Cell culture and generating the in vitro NAFLD huh7 model

Huh7 cells were selected to generate the steatotic *in vitro* hepatic model as a widely used reference. For all experiments, Huh7 cells were cultured in DMEM media supplemented with fetal bovine serum (10 %) and antibiotics (penicillin, 100 U mL−1 and streptomycin, 100 μg mL−1) and maintained at 37 °C under 5 % CO_2_. The in vitro model of steatotic cells was created by incubating hepatic Huh7 cells with 200μM palmitic acid (PA) BSA-NaOH solution, which induces a similar morphology to hepatic steatosis (De Gottardi et al., 2007; Gómez-Lechón et al., 2007). Fatty acid (FA) solution was made by dissolving PA in 0.1M NaOH to 0.1M PA, then combining this with 10% FA-free BSA in DMEM solution to a concentration of 10mM PA-BSA to obtain a molar ratio FA:BSA of 6.67:1. Huh7 cells were seeded in 6 well plates at a density of 0.5×106 cells per well the day before fatty acid treatment. Next day, FA-conjugate was diluted to 200uM final concentration in the culture medium and incubated with cells for 24 hours. This protocol was modified from previously described protocols (Listenberger et al., 2001). We evaluated changes in lipid composition and cell morphology by lipid staining followed by microscopy (Figure S5). We measured the impact of the steatosis on fetuin-B secretion with quantitative WB (Figure S6).

### RNA-Seq of NAFLD cell lines

RNA extraction for non-treated and PA-treated Huh7 was performed in triplicate. Cells were lysed in culture flask using TRI reagent (Millipore Sigma), and total RNA were extracted using Direct-zol™ RNA Microprep (Zymo Research) according to the manufacture instructions. RNA was quantified using NanoDrop according to the manufacturer’s instructions. The RNA integrity number (RIN) for each sample was measured using an Agilent 2200 TapeStation. Samples were sent to The UCSD IGM Genomics Center for mRNA library Prep and sequencing. Total RNA was assessed for quality using an Agilent Tapestation 4200, and samples with an RNA Integrity Number (RIN) greater than 8.0 were used to generate RNA sequencing libraries using the TruSeq Stranded mRNA Sample Prep Kit with TruSeq Unique Dual Indexes (Illumina, San Diego, CA). Samples were processed following manufacturer’s instructions, starting with 1 ug of RNA and modifying RNA shear time to five minutes. Resulting libraries were multiplexed and sequenced with 100 basepair (bp) paired end reads (PE100) to a depth of approximately 25 million reads per sample on an Illumina NovaSeq 6000. Samples were demultiplexed using bcl2fastq v2.20 Conversion Software (Illumina, San Diego, CA). Sequence data for RNA-Seq were quality controlled using FastQC and summarized with multiQC (Ewels et al., 2016). Low-quality bases were trimmed from the reads with Trimmomatic (Bolger et al., 2014). Reads were then mapped to the human GRCh38.p12 genome and quantified with Salmon (Patro et al., 2017).

### CellFie analysis comparing WT vs NAFLD model cells

Expression data were run through secCellFie using the secretory expansion of genome-scale model Recon2.2 (Swainston et al., 2016) with the following parameters: local mean threshold with upper and lower percentiles of 25 and 75 respectively. Hierarchical clustering of tasks scores was performed and visualized using the pheatmap package (Kolde, 2019) in R. Hypergeometric enrichment of CellFie systems in each cluster was carried out using the clusterProfiler package (Yu et al., 2012) in R. Differentially active tasks between WT and NAFLD model cells were calculated using Welch’s t-test. Significant tasks (|FC| ≥ 1.25, FDR < 0.1) were visualized in a volcano plot using the EnhancedVolcano package (Blighe et al., 2021) in R. To contextualize the fetuin-B interactome in terms of the globally altered secretory task, we first extracted the list of genes involved in each task using the GPR rules associated with the essential reactions. These genes were filtered to only include those with recorded interactions with fetuin-B, and this network of tasks and genes was visualized using ggraph package (Pedersen, 2022) in R. Edges of the graph were colored according to interaction strength with fetuin-B, and nodes colored according to gene LFC.

### Biotinylation by antibody recognition (BAR)

BAR was applied to triplicate samples of the fixed PA-treated and control Huh7 cells, targeting the product of fetuin-B to identify its PPIs. Briefly, cells were fixed with 4.0% paraformaldehyde (PFA) in PBST for ten minutes and then permeabilized with 0.4% PBST for ten minutes. Following this, peroxidases were inactivated with 0.2% H_2_O_2_ in PBS for ten minutes and then blocked for two hours with 5.0% normal goat serum in PBST. Cells were then incubated overnight at 4°C with rabbit anti-FetiunB antibody (1:100, 18052-1-AP), followed by a one hour intubation with anti-rabbit HRP-conjugated antibody (1:1000, ab7090). Proximal protein biotinylation occurs with treatment of H_2_O_2_ and tyramide-biotin (TSA Biotin System, Akoya Bioscience NEL700A001KT), resulting in tyramide-biotin radicalization and covalent deposition onto proximal proteins. Tyramide biotin stock solution was added to each sample and incubated for 5 minutes before addition of the H_2_O_2_ Amplification Diluent at the manufacturers’ recommended ratio of 1:50, tyramide to diluent. The reaction proceeded for three minutes before being stopped with 500 mM Sodium Ascorbate solution in PBST. To ensure expected localization of the biotinylation to the endomembrane system, a subsample was stained by dyed antibody and streptavidin and analyzed by confocal microscopy. Biotinylated proteins were pulled down from cell lysate using a streptavidin column, and then subjected to LC-MS/MS label free proteomic quantification (Clough et al., 2009). This way, the BAR method will quantify the changes in PPIs in between control and steatotic cells.

### Mass Spectrometry of WT and NAFLD cell lines

Cells were grown in 225 cm^2^ flask (one plate per biological replicate in triplicate) to approximately 70% confluence in complete media. Cells were labeled by BAR, (see above) harvested, and were lysed by boiling in a lysis buffer containing 2% SDS. Extracted proteins were centrifuged at 14,000xg to remove cellular debris and quantified by BCA assay (Thermo Scientific) as per manufacturer recommendations. Affinity purification of biotinylated proteins was carried out in a Bravo AssayMap platform (Agilent) using AssayMap streptavidin cartridges (Agilent), and the bound proteins were subjected to on-cartridge digestion with mass spec grade Trypsin/Lys-C Rapid digestion enzyme (Promega, Madison, WI) at 70°C for 2h. Digested peptides were then desalted in the Bravo platform using AssayMap C18 cartridges and the organic solvent was removed in a SpeedVac concentrator prior to LC-MS/MS analysis. Dried peptides were reconstituted with 2% acetonitrile, 0.1% formic acid, and analyzed by LC-MS/MS using a Proxeon EASY nanoLC system (Thermo Fisher Scientific) coupled to a Q-Exactive Plus mass spectrometer (Thermo Fisher Scientific). Peptides were separated using an analytical C18 Acclaim PepMap column 0.075 × 500 mm, 2µm particles (Thermo Scientific) in a 93-min linear gradient of 2-28% solvent B at a flow rate of 300nL/min. The mass spectrometer was operated in positive data-dependent acquisition mode. MS1 spectra were measured with a resolution of 70,000, an AGC target of 1e6 and a mass range from 350 to 1700 m/z. Up to 12 MS2 spectra per duty cycle were triggered, fragmented by HCD, and acquired with a resolution of 17,500 and an AGC target of 5e4, an isolation window of 1.6 m/z and a normalized collision energy of 25. Dynamic exclusion was enabled with a duration of 20 sec.

### MS data Analysis of WT and NAFLD cell lines

All mass spectra were analyzed with MaxQuant software (Tyanova et al., 2016) version 1.5.5.1. MS/MS spectra were searched against the *Homo sapiens* Uniprot protein sequence database (version January 2018) and GPM cRAP sequences (commonly known protein contaminants). Precursor mass tolerance was set to 20ppm and 4.5ppm for the first search where initial mass recalibration was completed and for the main search, respectively. Product ions were searched with a mass tolerance 0.5 Da. The maximum precursor ion charge state used for searching was 7. Carbamidomethylation of cysteines was searched as a fixed modification, while oxidation of methionines and acetylation of protein N-terminal were searched as variable modifications. Enzyme was set to trypsin in a specific mode and a maximum of two missed cleavages was allowed for searching. The target-decoy-based false discovery rate (FDR) filter for spectrum and protein identification was set to 1%. Enrichment of proteins in streptavidin affinity purifications were calculated as the ratio of intensity. To remove the systematic biases introduced during various steps of sample processing and data generation were normalized using the MaxQuant LFQ algorithm. The Proteus package (Gierlinski et al.,) was used to visualize the total protein number detected by mass spectrometry in each sample and perform principal component analysis (PCA). Perseus software (Tyanova and Cox, 2018) was employed for data preparation, filtering, and computation of differential protein abundance. The DEP package (Zhang et al., 2018) was used to explore whether missing values in the dataset are biased to lower intense proteins. Left-censored imputation was performed using random draws from shifted distribution. A Welch’s t-test with a multi-sample permutation-based correction for an FDR of 0.05 and S0 of 2 was employed to identify differentially expressed proteins using log2 transformed data.

## Results & Discussion

### Secretory expansion of CellFie predicts the activity of 21 core secretory pathway tasks

The concept of pathway “tasks’’ is not new and has been used to evaluate the quality and capabilities of genome-scale metabolic models (Agren et al., 2014; Blais et al., 2017; Bordbar et al., 2012; Duarte et al., 2007; Gille et al., 2010; Mardinoglu et al., 2014; Richelle et al., 2019a; Thiele et al., 2013). A task can be considered as a set of reactions that work together to accomplish a specific function (e.g., conversion of aspartate to arginine, synthesis of lactose, Krebs cycle - NADH generation). Recently, this concept was extended beyond a simple model benchmarking tool to quantify metabolic task activity using omic data (Masson et al., 2023; Richelle et al., 2021). This computational framework, dubbed CellFie, harnesses the mechanistic links in genome-scale metabolic models to predict the activity of hundreds of metabolic functions from omic data. Here, we use the genome-scale models of mammalian protein secretion (Gutierrez et al., 2020) to expand the tool to include tasks encompassing the core processes of protein secretion.

The secretory pathway consists of a complex network of molecular machinery responsible for the biosynthesis, processing, sorting, and delivery of both secreted and membrane proteins. Using species-specific genome-scale reconstructions of the secretory pathway (Gutierrez et al., 2020), we manually curated 21 secretory tasks for human, mouse, and Chinese hamster ovary cells covering the core processes involved in the synthesis of secreted and membrane proteins. Like the original metabolic tasks, the secretory tasks fall within a hierarchy of systems and subsystems covering five major secretory processes of the cell: i) translocation, ii) processing in the ER, iii) proteostasis, iv) processing in the Golgi, and v) vesicle trafficking (Figure 1). These additional tasks have been integrated with the CellFie framework, which can now be used to predict the metabolic and secretory capability of cells and tissues using omic data.

**Figure 1.**
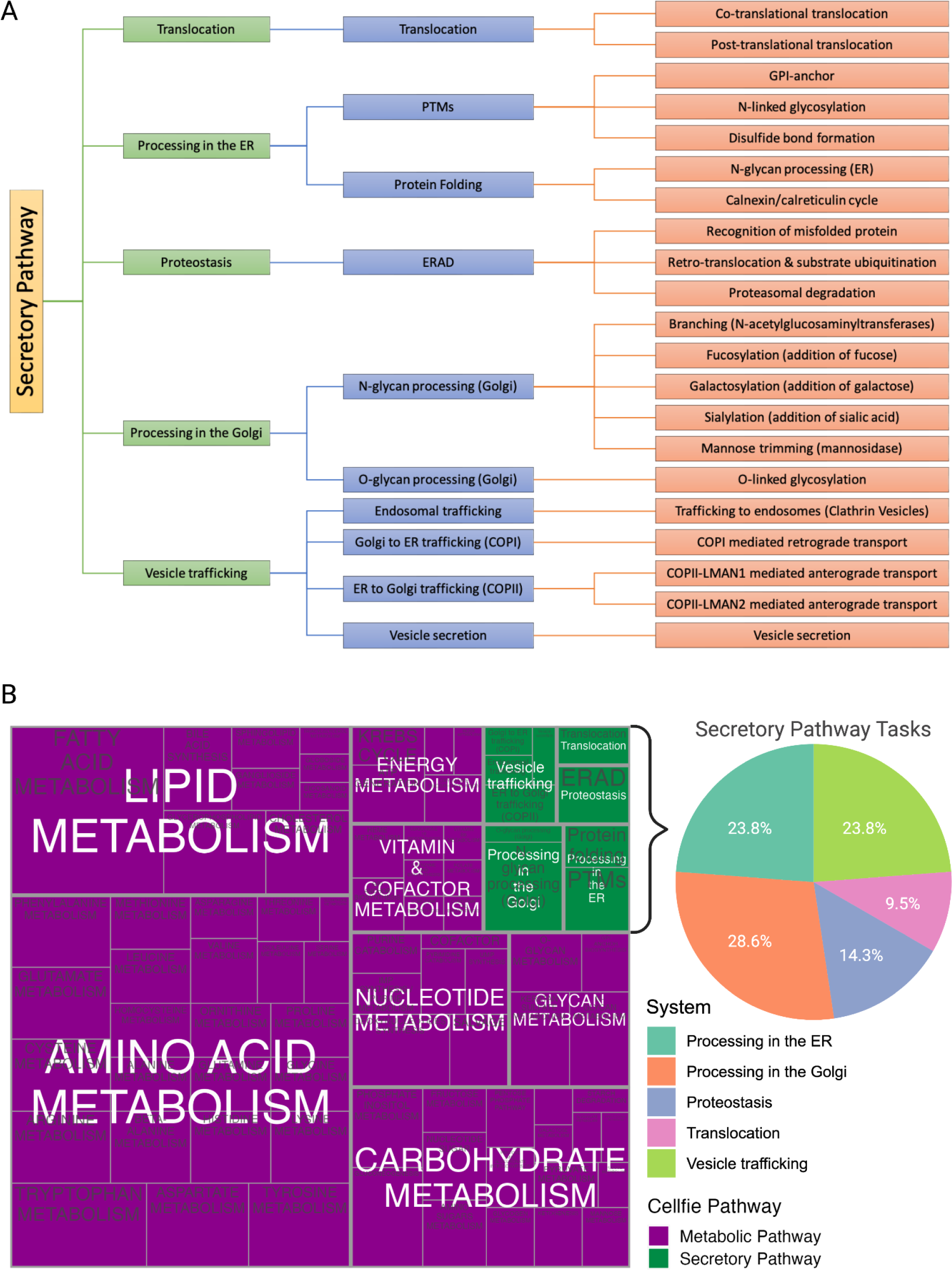
Integration of the secretory pathway with CellFie. **A)** Hierarchical organization of secretory systems (5), subsystems (10), and tasks (21) integrated with CellFie. **B)** Treemap depicting the hierarchical organization of CellFie systems and subsystems. Rectangle size is proportional to the number of tasks belonging to each subsystem, which have been colored according to pathway. The pie chart to the right highlights the addition of new secretory tasks and systems.

### Addition of secretory tasks increases clustering accuracy of immune cells

Immune cells originate from hematopoietic stem cells and differentiate into functionally-specialized cells that are distributed throughout the body to provide rapid immune response. Here, we used secCellFie to systematically assess the metabolic and secretory signatures differentiating immune cell types and subtypes with transcriptomic data from HPA (Uhlen et al., 2019) (Figure 2A). Hierarchical clustering of task scores using the original version of CellFie containing only metabolic information was unable to cleanly distinguish several of the immune cell groups including NK-cells and the T-cells (Figure 2B, top). Upon integration of secretory task information with the secCellFie, we were able to recover clusters of the immune cell types (Figure 2B, bottom). For the immune cell types that did not cluster together, the dendritic cells (DCs), we find that the individual clusters correspond to cell subtypes. To quantify how much secretory task information boosted clustering capabilities, we compared the resulting CellFie and secCellFie dendrograms with the dendrogram representing known immune cell lineage using the Fowlkes-Mallows Index (FM-Index) correlation. We found that integration of secretory tasks boosted correlation from 0.59 to 0.70 (Figure 2D). Thus, while the secretory pathway accounts for only a small fraction of the overall tasks in secCellFie, the addition of secretory task information improves functional characterization of cells and ultimately enables greater resolution to distinguish immune cell types.

**Figure 2.**
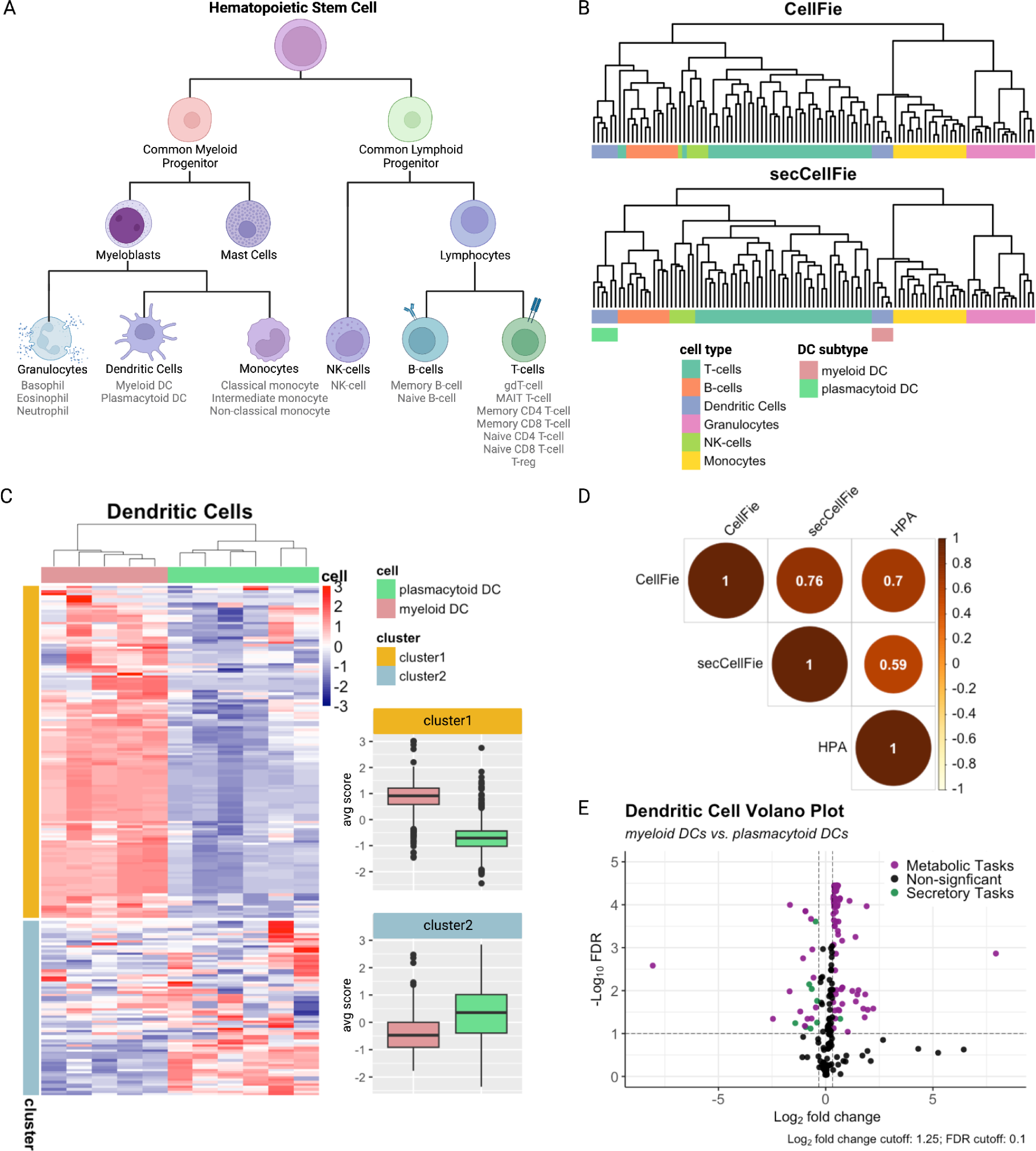
CellFie analysis using immune cell transcriptomic data from HPA. **A)** The Human Protein Atlas (HPA) immune cell data used in this study includes genome-wide RNA expression profiles for 6 single cell immune cell types and 18 subtypes which have been grouped based on main cell lineage. **B)** Hierarchical clustering of immune cell activity scores using CellFie, which only includes metabolic tasks (top) and the secretory expansion of CellFie, secCellFie (bottom). Cell types are indicated with distinct colors. **C)** Clustered heatmap of DC subtype activity scores. Boxplots to the right depict mean score within each cluster according to cell subtype. **D)** Correlation matrix between cell lineage dendrogram (HPA), dendrogram generated from hierarchical clustering of metabolic task scores (CellFie), and dendrogram generated from hierarchical clustering of metabolic and secretory scores (secCellFie). Area and color of the circles are proportional to the FM-Index correlation coefficient. The legend on the right shows the FM-Index correlation coefficients with their corresponding color. **E)** Volcano plot of CellFie scores comparing myeloid DCs vs. plasmacytoid DCs. Positive fold-changes denote higher activity in myeloid DCs and vice versa. Significant tasks are colored in pink and green according to the CellFie pathway.

### secCellFie captures cell-type specific immunological functions

We further sought to determine whether secCellFie could identify functional differences between subtypes of cells within the same cell type. To this end, we compared task scores of the two DC subtypes, myeloid DCs and plasmacytoid DCs. DCs represent a heterogeneous family of antigen presenting immune cells essential in adaptive immunity. While all DCs are capable of antigen uptake, processing, and presentation, DC subtypes have distinct markers and differ in immunological functions. Myeloid DCs are primarily involved in antigen presentation to T-cells to induce activation and effector differentiation, while plasmacytoid DCs are primarily involved in antiviral immune response and are characterized by high levels of type I interferons (INF-I) secretion.

The functional differences between these subtypes of DCs was clearly reflected in their unique metabolic and secretory signatures. Using hierarchical clustering, we identified two clusters of tasks with distinct trends (Figure 2C). Cluster 1 showed a general increase in myeloid DCs compared to plasmacytoid DCs, and was significantly enriched with metabolic tasks (FDR=6.2e-08). On the other hand, cluster 2 showed a general increase in plasmacytoid DCs compared to myeloid DCs, and was significantly enriched with secretory pathway tasks (FDR=6.2e-08). Using Welch’s t-test, we found that 8 of the 9 significantly differentially activated secretory tasks showed increased activity in plasmacytoid DCs (Figure 2E), consistent with the cluster observations. We hypothesize this is likely due to the massive production and secretion of INF-I observed in this subset of cells. Interestingly, the single secretory task with significantly greater activity in myeloid DCs was the GPI-anchor task, which involves the post-translational modification of proteins with glycosylphosphatidylinositol (GPI) to facilitate anchoring to the plasma membrane. We surmise this might be the result of increased membrane surface proteins used for antigen presentation characterizing this subset of DCs.

### secCellFie captures clone- and phase-specific signatures in recombinant protein-producing CHO cells

Recombinant protein (rProtein) production imposes considerable metabolic and secretory demand on the host cell. The metabolic burden of expressing rProteins can hinder cell growth and consequently result in overall lower biomass (Chusainow et al., 2009; Jiang et al., 2006; Ong et al., 2019; Pilbrough et al., 2009; Wu et al., 2016; Wurm, 2004). Additionally, competition for secretory pathway resources in rProtein synthesis imposes a significant burden on the secretory machinery of cells, which can result in lower expression of energetically expensive endogenous proteins (Gutierrez et al., 2020). To systematically assess the metabolic and secretory burden of rProtein in CHO cells, the dominant expression system used in biopharmaceutical production, we used secCellFie to analyze RNA-Seq data from high (A1) and low (A0) monoclonal antibody-producing CHO cells (Dahodwala et al., 2019; Jiang et al., 2006; Jiang and Sharfstein, 2009). Briefly, the high producing clone A1 represents the dihydrofolate reductase (DHRF)/methotrexate (MTX)-amplified progeny of parental cell line A0, and samples were taken in both exponential and stationary phase of batch shake-flask cultures (Figure 3A).

**Figure 3.**
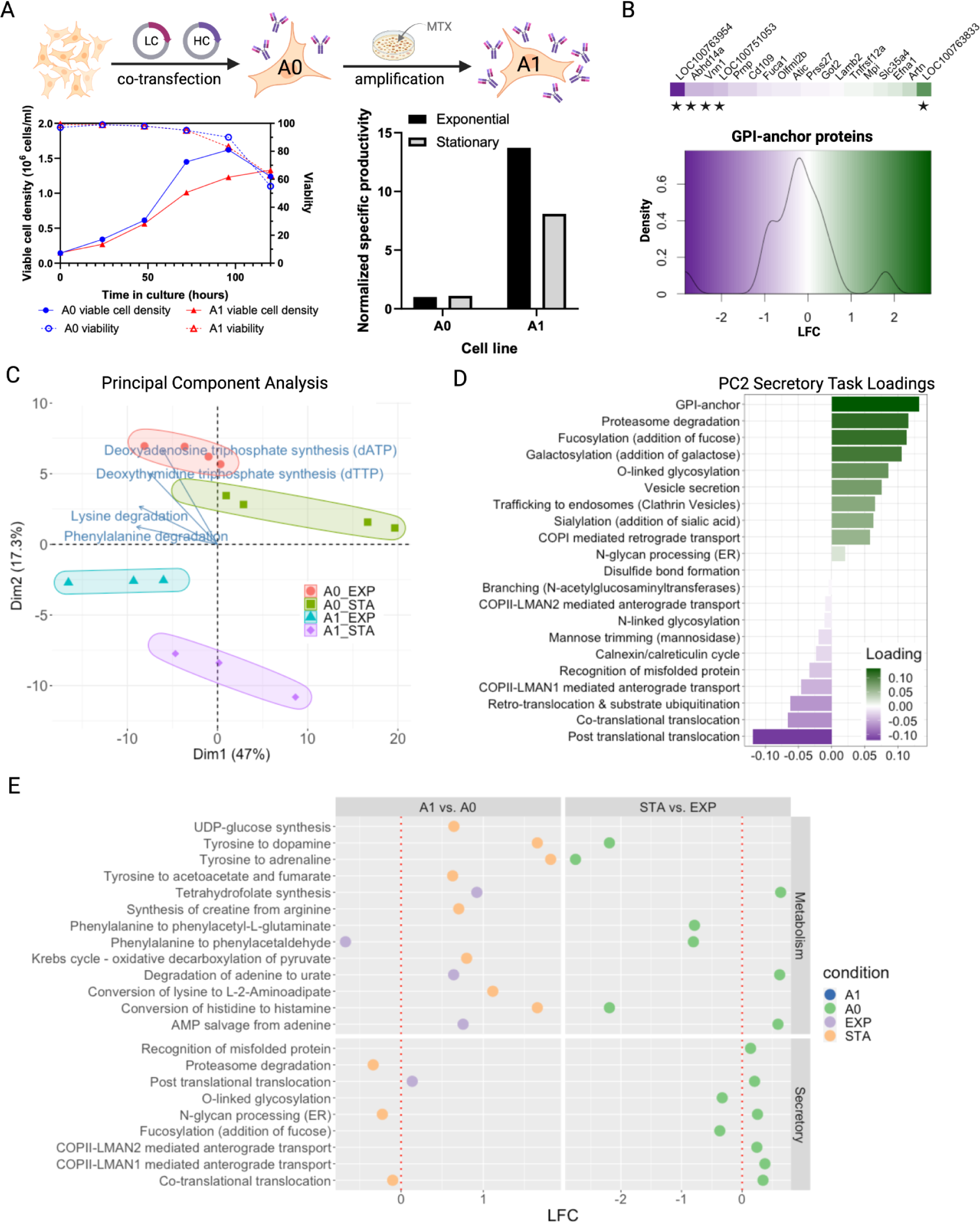
CellFie analysis on rProtein producing CHO cells. **A)** Schematic outlining cell line generation and experimental setup used in the study. DG-44 CHO cells were co-transfected with light- and heavy-chain containing plasmids. Gene amplification was performed by stepwise selection with increasing MTX concentrations. For this study, a low-producing parental cell line (A0) and its amplified high-producing progeny cell line (A1) were chosen for RNA-Seq analysis in both exponential and stationary phase. Figure adapted from (Dahodwala et al., 2019). **B)** Heatmap bar and density plot of log fold change (LFC) for genes encoding GPI-anchor proteins between A1 and A0 cells. Negative fold-change denotes lower expression in A1 and vice versa. Genes with statistically significant changes (LFC>= 1.5, FDR<=0.01) are indicated with a star. **C)** Biplot graph showing PCA score and top four loadings of CellFie tasks for A1 and A0 cells in exponential and stationary phase. **D)** Barplot showing the secretory pathway task vector loadings contributing to PC2. Colors represent the loading contribution to PC2. **E)** Dotplots of the significant differentially activated metabolic (top) and secretory (bottom) tasks between phases (right) and cell lines (left). A positive log fold change (LFC) between phases indicates higher activity in exponential phase and vice versa. Likewise, a positive LFC between cell lines denotes higher activity in A1 cells and vice versa.

Principal component analysis (PCA) using the task scores resulted in four distinct clusters representing each cell line in their respective phase (Figure 3C). For both the high and low producing clones, we find that samples collected in exponential phase are shifted up and to the left compared to their stationary phase counterparts. Coincidentally, the tasks with the strongest loadings show similar directionality. These tasks include synthesis of nucleotide base pairs adenine (A) and thymine (T), and degradation of the amino acids lysine and phenylalanine. Elevated DNA synthesis activity characterizing the clones in exponential phase is likely a reflection of increased cell replication and biomass production associated with exponential phase cell growth. Furthermore, the upregulation of lysine and phenylalanine degradation pathways, which result in nutrient and energy precursors including acetyl-CoA and acetoacetate, could be a result of diverting amino acid resources to fuel cellular replication.

The second principal component (PC2) clearly separated the high producing amplified cell line (A1) from the low producing parental cell line (A0). To understand how secretory processes contribute to this separation, we ranked the secretory tasks by their loadings on PC2 (Figure 3D). Synthesis of GPI-anchor proteins had the strongest effect on PC2, with elevated activity characterizing the low-producing clones. Competition for secretory pathway resources in rProtein synthesis imposes a considerable burden to the secretory machinery of the cells, which can result in lower expression of energetically expensive endogenous proteins in CHO (Gutierrez et al., 2020). These results could suggest that the high-producing clones are downregulating the production of energetically expensive GPI-anchor proteins to accommodate secretory requirements for high rProtein production. In fact, when we compare the expression level of genes encoding GPI-anchor proteins between A1 and A0, we find that A1 has indeed considerably downregulated the expression of several energetically expensive GPI-anchor proteins (Figure 3B). Additionally, we observe that post-translational translocation (characteristic of A1) and proteasomal degradation (characteristic of A0) represent the next two loading vectors with the largest effect on PC2. This could be further contributing to the higher production of rProtein compared to A0.

### secCellFie identifies shift in activity that could explain productivity phenotypes

To identify changes in metabolic and secretory activity, we performed a pairwise comparison of task scores between cell lines and phases (Figure 3E). We observed considerable shifts in secretory activity between phases in the A0 cell line, while A1 showed no significant changes between phases (bottom right). While cell lines showed minor secretory differences in exponential phase, we found that as cells shift into stationary phase, A0 cells significantly increased several secretory pathway tasks compared to A1 (bottom left) including translocation (FDR=0.08), N-glycan processing in the ER (FDR=0.07), and proteasomal degradation (FDR=0.07). Overall, these results suggest that the high producing cell line A1 is able to maintain efficient protein synthesis throughout phases, while the low-producing parental cell line A0 is potentially overwhelmed with processing in the ER from increased translocation in stationary phase, ultimately resulting in greater protein clearance and low titers. Measurements confirm that the specific productivity remains significantly higher in A1 cells compared to A0 throughout phases.

Similarly, we found no significant metabolic changes in A1 as it transitioned from exponential to stationary phase, while many metabolic shifts were observed in A0 (Figure 3E, top right). This could again be indicative of a high-producing phenotype maintained across phases in A1. From a metabolic standpoint, exponential phase is typically associated with high biomass production while stationary phase is characterized by nutrient depletion due to high protein production. However, here we found significant decreases in nutrient-depleting amino acid degradation tasks – i.e., tyrosine to adrenaline (FDR=0.01), conversion of histidine to histamine (FDR=0.02), tyrosine to dopamine (0.02), phenylalanine to phenylacetaldehyde (FDR=0.03), and phenylalanine to phenylacetyl-L-glutaminate (FDR=0.03) – in A0 stationary phase. Additionally, A0 cells trended toward a significant decrease (FDR=0.06) in tetrahydrofolate (THF) synthesis (biomass component in metabolic model) in exponential phase. These A0 metabolic signatures oppose typical phase characteristics. Interestingly, many of the same amino acid degradation tasks showed considerably greater activity during stationary phase in A1 cells compared to A0 (top left). A1 cells also exhibited increases (FDR=0.09) in THF synthesis compared to A0 in exponential phase, however this is most likely an artifact of the amplification process using DHFR which functions in the THF synthesis pathway. Furthermore, we found A1 cells significantly increased (FDR=0.04) UDP-glucose synthesis compared to A0 in stationary phase. UDP-glucose is the starting point for glycosylation, a critical quality attribute of mAb therapeutics. Altogether, the distinct metabolic signatures identified here may explain the higher productivity phenotype of A1, providing hypotheses for future studies.

### secCellFie identifies altered metabolic and secretory function in an in vitro model of NAFLD

The liver regulates glucose and lipid metabolism by secreting hepatokine hormones into the blood. Consequently, perturbed hepatokine secretion is associated with many metabolic disorders including nonalcoholic fatty liver disease (NAFLD) and non-alcoholic steatohepatitis (NASH). Hepatic steatosis in NAFLD and NASH changes the secretion profile of approximately 20% of classic hepatokines (Meex et al., 2015), and these changes in hepatokine secretion can contribute to the pathology of metabolic disease by disrupting systemic metabolism and energy homeostasis, leading to dysregulated gluconeogenesis, dyslipidemia, and pathogenesis (Lebensztejn et al., 2016; Meex and Watt, 2017). However, changes in hepatokine secretion in steatotic hepatocytes often do not correlate with differential mRNA expression (Fu et al., 2012; Gorden et al., 2015), implicating the secretory pathway as a likely source of the perturbed secretion. To study the metabolic and secretory alterations in hepatic steatosis, we developed an in vitro model of NAFLD. Briefly, Huh7 cells were treated with palmitic acid (PA) to induce fat accumulation mimicking hepatic steatosis. Changes in lipid composition and cell morphology were evaluated using lipid staining and microscopy (Figure S5).

To systematically characterize the global metabolic and secretory dysregulation in NAFLD, we performed a CellFie analysis on RNA-Seq from normal (WT) and PA-treated NAFLD model cells. Hierarchical clustering of secCellFie tasks showed distinct metabolic and secretory fingerprints of NAFLD (Figure 4A). Cluster A, which showed increased activity in NAFLD model cells compared to WT, was significantly enriched (FDR=2.96e-05) with tasks involved in carbohydrates and lipid metabolism (FDR=7.38e-03). In particular fatty acid metabolism was significantly enriched (FDR=3.96e-04) in Cluster A. This is unsurprising given that NAFLD model cells were treated with the fatty acid PA to induce steatosis, and alterations in lipid metabolism and abnormal lipid accumulation, especially fatty acids, are a hallmark of NAFLD (Hliwa et al., 2021; Pei et al., 2020). On the other hand, Cluster B, which showed limited activity in NAFLD model cells compared to WT, was significantly enriched with tasks involved in amino acid (FDR=0.01) and nucleotide metabolism (FDR=0.03). Several studies report altered amino acid levels in hepatic steatosis (van den Berg et al., 2019; Gaggini et al., 2018; Hasegawa et al., 2020; Lake et al., 2015; Mardinoglu et al., 2017). Consistent with our results, they hypothesize that mitochondrial dysfunction in NAFLD could result in impaired amino acid metabolism (van den Berg et al., 2019).

**Figure 4.**
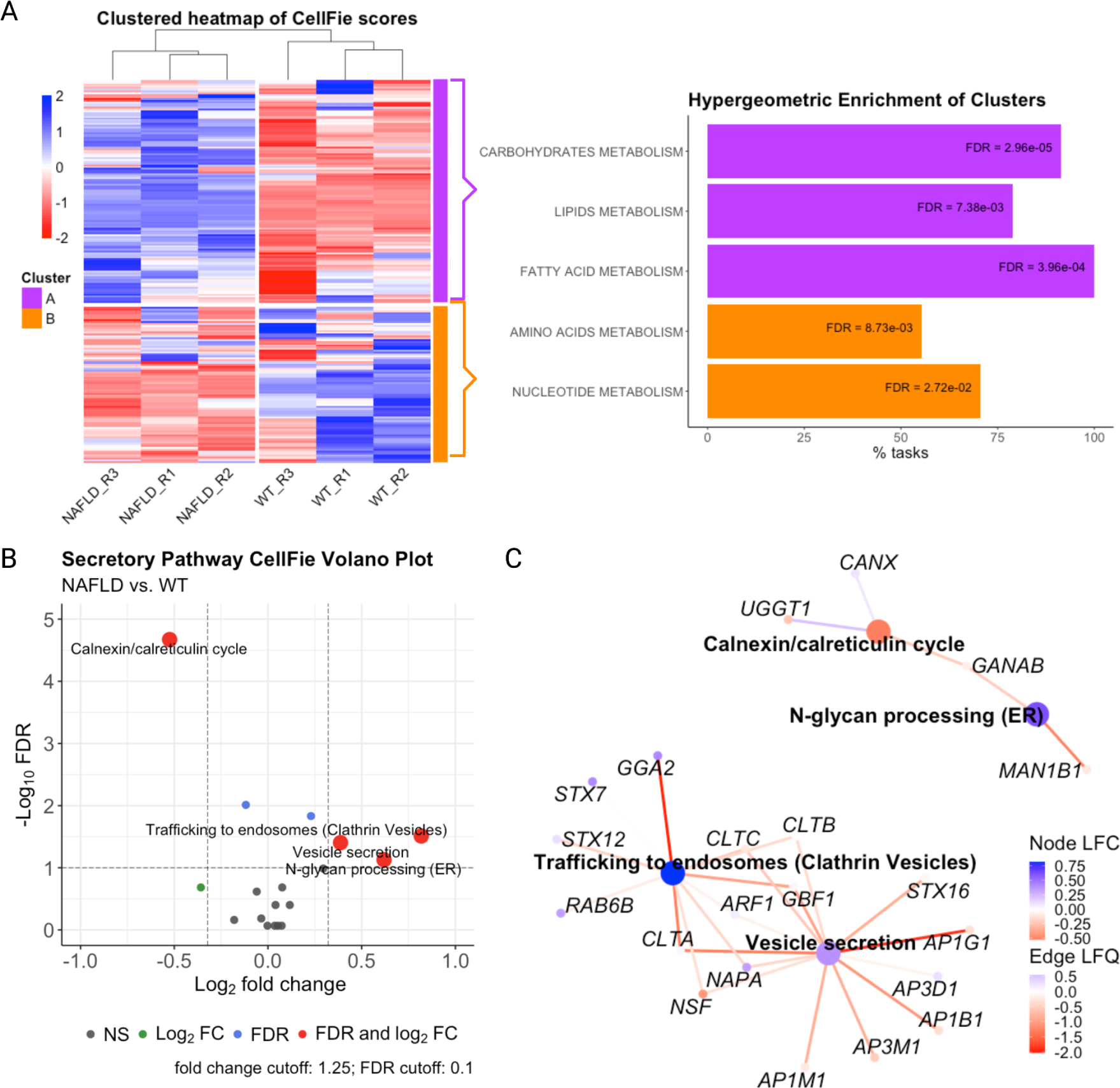
CellFie analysis in an NAFLD cell model. **A)** Clustered heatmap of CellFie scores comparing NAFLD model and WT cells. Barplot to the right shows hypergeometric enrichment analysis of significantly enriched CellFie systems and subsystems for each cluster. The x-axis corresponds to the percentage of tasks from the system or subsystem present in the cluster. **B)** Volcano plot of secretory task scores comparing NAFLD model cells vs. WT. Positive fold-changes denote higher activity in NAFLD calls and vice-versa. Tasks that passed both significance (FDR) and fold change thresholds are shown in red. **C)** Network diagram of the differentially activated secretory tasks and contributing genes that has been overlaid with the fetuin-B interactome. Nodes have been colored according to LFC in either expression (gene) or activity (task). The network has been filtered to only include proteins that interact with fetuin-B with edges colored according to interaction strength (LFQ) with fetuin-B. Positive fold-changes in node LFC and interaction strength LFQ denote greater expression/activity/interaction strength in NAFLD model cells, and vice-versa.

To further understand the alterations of secretory functions during hepatic steatosis, we performed differential activation analysis on the secretory task scores (Figure 4B). The calnexin/calreticulin cycle, a chaperone-mediated system for glycoprotein folding and quality control in the ER, was the only task that showed significantly lower (FDR=2.49e-05) activity in NAFLD compared to WT. We identified three other tasks with significantly higher activity in NAFLD compared to WT: N-glycan processing in the ER (FDR=8.69e-02), clathrin-coated vesicle trafficking to endosomes (FDR=3.59e-02), and vesicle secretion (FDR=4.61e-02). Previous studies have shown that lipid accumulation in hepatocytes stimulates the release of extracellular vesicles (Akazawa and Nakao, 2018; Hirsova et al., 2016; Liao et al., 2018; Yuan et al., 2023), which is in accordance with the results observed here. Together these results indicate that steatotic cells transcriptionally downregulate chaperone-mediated protein folding, and upregulate protein-processing and post-Golgi vesicle trafficking capabilities.

### Contextualizing the interactome of model hepatokine fetuin-B within the secCellFie framework reveals potential drivers of hepatokine dysregulation in NAFLD

We were particularly interested in understanding how the global changes in cell function are associated with the dysregulated secretion of hepatokines. We chose to study the model hepatokine fetuin-B, which demonstrates elevated secretion without significant transcriptional changes (Meex et al., 2015) (Figure S6). To identify the cell machinery involved in fetuin-B synthesis and secretion, we implemented the Biotinylation by Antibody Recognition (BAR) assay. The BAR method is an antibody-guided proximity-biotinylation labeling approach that enables *in situ* detection of transient protein-protein interactions (PPIs) on any native bait protein (Bar et al., 2018) (Figure S8A). BAR successfully biotinylated proteins proximal to our bait protein, fetuin-B (Figure S7); and using mass spectrometry we identified 1505 unique biotinylated proteins corresponding to the fetuin-B interactome (Supplementary Data 2).

To contextualize the globally altered CellFie tasks with the dysregulated secretion of fetuin-B in NAFLD, we overlaid the fetuin-B interactome on the network of genes involved in the significantly altered secretory tasks (Figure 4C). While steatotic cells were characterized by a global reduction in calnexin/calreticulin activity, we found greater interaction strength between fetuin-B and many of the proteins involved in that task. Additionally, fetuin-B displayed weaker interactions with N-glycan processing in the ER and post-Golgi trafficking secretory machinery proteins even though we observe a global increase in activity in those tasks. Altogether, while results indicate a global shift in secretory capacity at the transcriptional level, fetuin-B-specific interactions show an opposing pattern with the secretory machinery proteins involved in these altered tasks.

We hypothesize two possible mechanisms that could explain this interesting discord: i) the resulting stressors of hepatic steatosis induce unconventional secretion of fetuin-B via the Golgi-bypass route (Balmer and Faso, 2021; Grieve and Rabouille, 2011; Rabouille, 2017), or ii) the global increase in post-Golgi trafficking genes induced by NAFLD generate an excess of available secretory machinery and thus alleviate vesicle trafficking bottlenecks in fetuin-B secretion. To test the Golgi-bypass theory, we conducted a co-localization experiment between fetuin-B and the Golgi-localized protein Golgi SNAP Receptor Complex Member 1 (GosR1), an essential component of the Golgi SNAP receptor (SNARE) complex (Figure S9A). We found no significant changes in co-localization between fetuin-B and GosR1 in PA-treated vs WT cells (Figure S9B), concluding that fetuin-B is likely not secreted via Golgi-bypass mechanisms. We therefore propose that the global increase in vesicle secretion induced in hepatic steatosis alleviates vesicle trafficking bottlenecks. Under WT conditions fetuin-B is accumulating at the vesicle trafficking step and forming an increasing number of interactions while waiting in the “secretory queue”. The global increase in vesicle secretion machinery alleviates this queue, as seen by the reduced number of interactions of fetuin-B with vesicle secretion machinery proteins, thus leading to greater secretion without significant increases in mRNA expression. Additional studies will be needed to validate this hypothesis. Ultimately, a fundamental understanding of how hepatokine synthesis and secretion is regulated is essential to efforts seeking to diagnose and treat disorders such as NAFLD, and tools such as CellFie can generate novel hypotheses of NAFLD-associated disease mechanisms.

## Conclusion

Enrichment analyses using omic data, such as GSEA and hypergeometric enrichment, are invaluable tools commonly used in the scientific community (Khatri et al., 2012) to identify sets of genes and proteins that are significantly over- or under-represented in a sample. However, to obtain a more mechanistic understanding of the molecular changes underlying phenotype, we developed the CellFie tool which leverages the mechanistic relationships provided by genome-scale models. Here, we expand the scope of CellFie beyond metabolism to include 21 novel secretory modules. Using a wide range of datasets, we highlighted the power of this tool to gain phenotype-relevant interpretation of complex data types and generate hypotheses in various contexts ranging from biomedical questions to biopharmaceutical manufacturing.

First, we demonstrate how the inclusion of secretory tasks with CellFie enhanced the tool’s ability to classify immune cells types and capture cell-type specific immunological functions. Then, we illustrate how secCellFie can be used to understand the molecular mechanisms underlying rProtein production and disease. The work here not only presents a novel tool for systematic analysis of secretory function, but also demonstrates novel implementation of the revolutionary BAR method to establish protein-specific secretory interactomes. The methods presented here can easily be leveraged in rProtein production to investigate protein-specific secretory machinery as potential targets for cell line engineering. Collectively, the series of vignettes demonstrate the value of this tool to monitor and quantify multiple complex biological systems providing phenotypic interpretation from omic data. This easy-to-use tool enables scientists, regardless of their computational background, to estimate the activities of metabolism, cytokine secretion, receptor production, and recombinant protein bioproduction.

## Acknowledgements

This work was supported by generous funding from NIGMS (R35 GM119850), NIAID (UH2AI153029 to N.E.L.), NSF (CBET-2030039), and the Novo Nordisk Foundation (NNF20SA0066621). This publication includes data generated at the UC San Diego IGM Genomics Center utilizing an Illumina NovaSeq 6000 that was purchased with funding from a National Institutes of Health SIG grant (#S10 OD026929).

## Author Contributions

H.O.M and N.E.L implemented the expansion, conducted the analyses, and wrote the paper. M.S., C.M.R., L.W, K.S. A.R.C, and C.C.K generated and processed the data used in the NAFLD vignette. V.T, H.D, P.M, B.G.V., and S.T.S generated and shared the data used in the rProtein vignette.

## Data Availability

The CellFie algorithm and code for the vignettes are available on [GitHub](https://github.com/LewisLabUCSD/CellFie_SecretoryPathway).

