## Supplementary Information for "Inferring secretory and metabolic pathway activity from omic data with secCellFie"

**This file contains:**

**Supplementary Results**

**Supplementary Methods**

**Supplementary Figures 1-9**

**Supplementary References**

**Other supplementary materials for this manuscript include the following:**

**Supplementary Data 1 –** Table of essential reactions for the 21 secretory tasks for CHO (sheet 1), mouse (sheet 2), and human (sheet 3).

**Supplementary Data 2 –** Fetuin-B interactome prepared using the Perseus software to filter and compute differential protein abundance.

### Supplementary Results

#### BAR successfully tracks altered fetuin-B secretion and shows condition specific patterns in our NAFLD model

To elucidate the source of changes in hepatokine secretion, we developed an *in vitro* model of NAFLD using Huh7 cells and studied the secretion of model hepatokine fetuin-B, which demonstrates elevated secretion without significant transcriptional changes (Supplementary Figure S6). We implemented the Biotinylation by Antibody Recognition (BAR) assay to identify the cell machinery involved in fetuin-B secretion. The BAR method is an antibody-guided proximity-biotinylation labeling approach that enables in situ detection of transient protein-protein interactions (PPIs) on any native bait protein (Bar et al., 2018) (Figure S8A). Here, we demonstrated that BAR successfully biotinylated proteins proximal to our bait protein, fetuin-B (Figure S7). Identified PPIs included proteins involved in the trafficking and processing of secreted proteins. We subsequently identified and quantified the biotinylated proteins in normal and NAFLD model cells to identify which protein interactions were altered. To do this the biotinylated proteins were labeled and subjected to mass spectrometry analysis. We identified 1505 unique biotinylated proteins corresponding to the fetuin-B interactome. Surprisingly, only 5 proteins showed significant differential interactions (FDR 0.1; LFQ 1.5) between normal and NAFLD model cells (Figure S8B).

Hierarchical clustering of the biotinylated proteins revealed condition-specific responses of the fetuin-B interactome. One of the samples, NAFLD_R2, appeared to be an outlier compared to the other FA-treated samples. We believe this may be the result of a batch issue caused by sample loss during the BAR experiment. Clustering of the interactome identified two distinct classes of proteins with altered interactions (Figure S8C). While the proteins in Cluster 1 show a general depletion of interactions with fetuin-B in our NAFLD model, the proteins in Cluster 2 show the opposite trend and tend to have increased interactions with fetuin-B in our NAFLD model. Hypergeometric enrichment revealed that the proteins in Cluster 1 are enriched in the mitochondrion, mitochondrial matrix, and Golgi membrane compartments. Meanwhile, Cluster 2 is enriched with proteins involved in unfolded protein binding, glycolysis/gluconeogenesis, antigen processing and presentation, and lipid and atherosclerosis.

### Supplementary Methods

#### Immunofluorescence NAFLD cell lines

Immunofluorescence staining was used to study cellular localization and lipid composition. For lipid composition staining, control and fatty acid treated cells were were stained in 12-well plates using LipidSpot 488 (Biotium) and 10 µg/ml DAPI for 30 minutes at 37°C. Plates were then imaged using confocal microscopy with fluorescence and brightfield for visualization of lipid droplet accumulation. For colocalization studies, a subsample of the BAR labeled cells were taken and probed with goat anti-rabbit-Dylight 650 conjugate (1:300, Thermofisher) and streptavidin-DyLight 594 conjugate (1:1000, Thermofisher) targeting the biotinylated proteins. For fetuin-B Golgi interactions, fixed control and PA-treated Huh7 cells were fixed and stained with rabbit anti-FetiunB antibody (1:100, 18052-1-AP) and mouse anti-GosR1 (1:100, ab88462), followed by goat anti-rabbit Alexa 488 (1:500, ab150077) and goat anti-mouse Alexa 647 (1:500, ab150115) secondary antibodies. Cells were then washed, counterstained with DAPI, mounted on the slide using antifade vectashield mountant, and imaged using Leica SP8 Confocal with Lightning Deconvolution. Colocalization quantification was performed for the deconvolved images using Fiji's (ImageJ 1.52p) Coloc_2 analysis tool (Schindelin et al., 2015). Pixel intensity colocalization of two channels was evaluated by Pearson's Coefficient (range: -1.0 to 1.0). Background pixel intensity was subtracted using Fiji's rolling ball algorithm and a region of interest (ROI). Thresholds were determined using Coloc_2's bisection method, which is further used to adjust for background. Above threshold metrics were reported.

#### Western blotting of NAFLD cell lines

Semi-quantitative WB was used to validate the impact of the lipid composition on fetuin-B expression and efficiency of biotinylation in the labeled cells. To measure impact of fatty acid treatment on fetuin-B secretion, control and PA-treated Huh7 cultures were prepared in triplicates using a 6-well plate. Cells taken were harvested and lysed using RIPA buffer. Protein content of the cell lysate was quantified using a Pierce BCA protein assay (Thermofisher). To measure the intracellular level of fetuin-B, 20 ug of total protein from the cell lysate and was loaded on SDS-PAGE gel for electrophoresis and resolved proteins were transblotted to nitrocellulose membrane using Trans-Blot Turbo Transfer System from Bio-RAD. To measure secreted fetuin-B, media sans FBS (which contains fetuin-B that can mask results) from Huh7 control and PA-treated cultures was centrifuged to remove cell debris and 22.5$\mu$l of the supernatant was loaded on a SDS-PAGE gel and transblotted in a similar fashion. The membranes were blocked with LiCOR Intercept (TBS) Blocking Buffer and probed with rabbit anti-fetuin-B antibody diluted at 1:1000, and additionally for the lysate membrane mouse anti-βactin (1:10,000, 66009-1-Ig), overnight at 4ºC. The membranes were then washed and incubated with LiCOR IRDye goat anti-mouse 680 and/or goat anti-rabbit 800 at 1:10000 for one hour. Membranes were imaged using the LiCOR Odyssey Imaging System. For staining of intracellular biotinylated proteins, 20 ug of total protein from BAR and control labeled Huh7 control and PA-treated lysates were loaded and resolved and transblotted as described above. For visualizing the proteins' bands, blocking reagents were the same as above, LiCOR IRDye streptavidin 680 at 1:1000 dilution was used, and the membranes were imaged with the same Odyssey system.

#### qRT-PCR of NAFLD cell lines

Triplicate cultures of control and PA-treated Huh7 cultures were grown in 6-well plates. RNA extraction for 1×10^6 cells from each well was performed (Direct-zol RNA MicroPrep, Zymo), RNA extracts were quantified by NanoDrop, and RT for cDNA synthesis was performed using 1000ng RNA (iScript, BioRad). Samples were run at 25ºC for 5 minutes, 46ºC for 20 minutes, 95ºC for 1 minute, and stored at 4ºC. For RT-qPCR, iTaq universal SYBR Green Supermix (BioRad) was used according to the manufacturer's instructions. Samples were run at 95ºC for 30 seconds followed by 40 cycles of 95ºC for 5 seconds and 53ºC for 25 seconds. PCR products were evaluated by melt curve and agarose gel electrophoresis. Primers used were fetuin-B (IDT) and reference genes RPS18 (IDT) and SRSF4 (IDT).

### Supplementary Figures

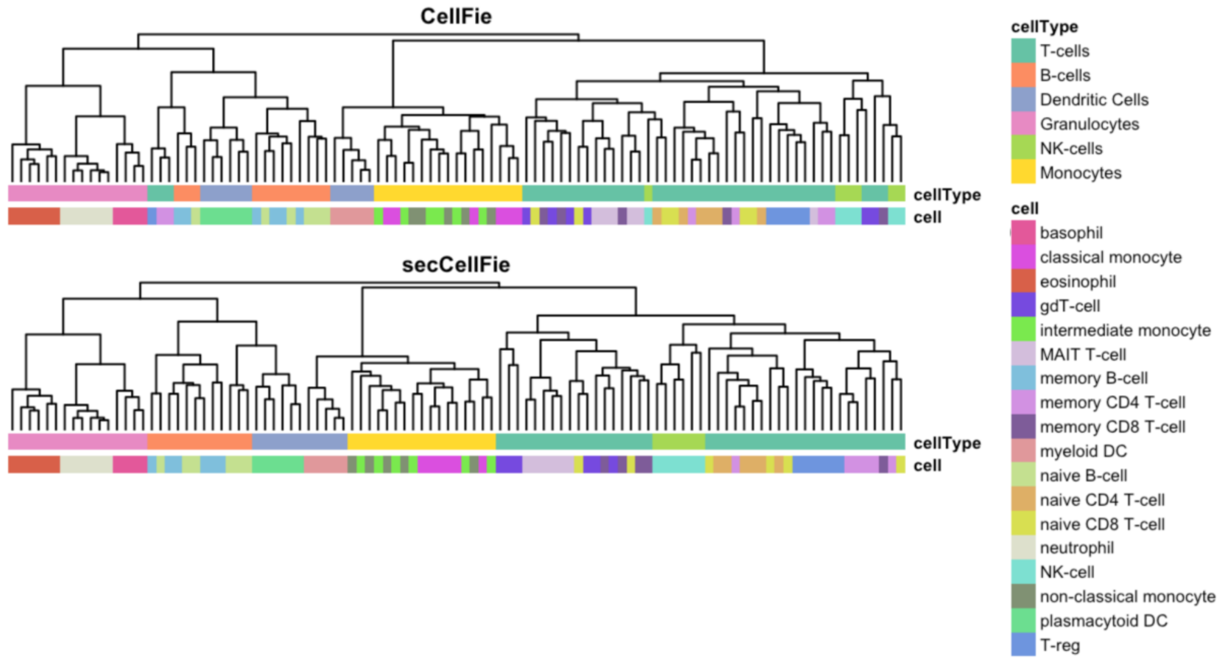


**Supplementary Figure S1**. Hierarchical clustering of immune cell CellFie scores using the old version of CellFie (top) and the secretory expansion of CellFie (bottom). Cell types (cellType) and subtypes (cell) are indicated using distinct colors.

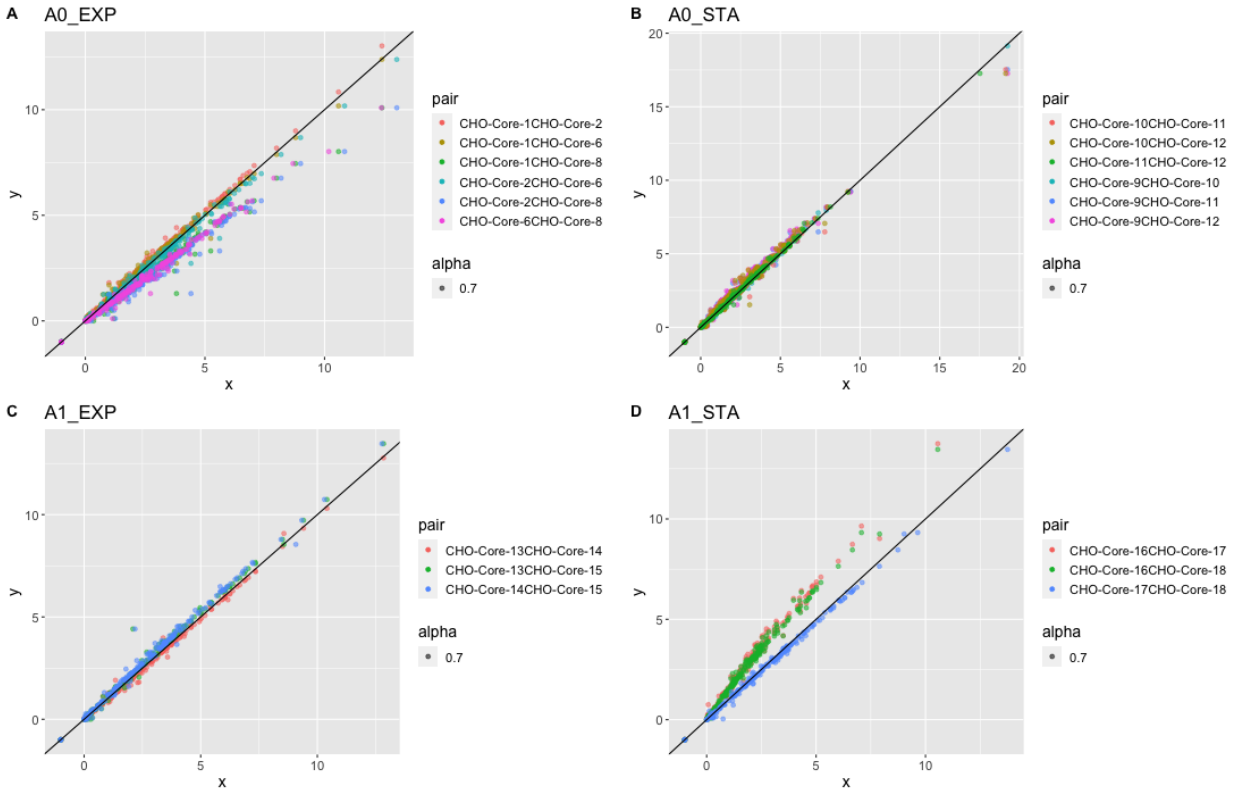


**Supplementary Figure S2**. CellFie scores of IgG producing CHO cell lines before normalization. You can see that CHO-Core-8 (A0_EXP) and CHO-Core-16 (A1_STA) appear to be skewed.

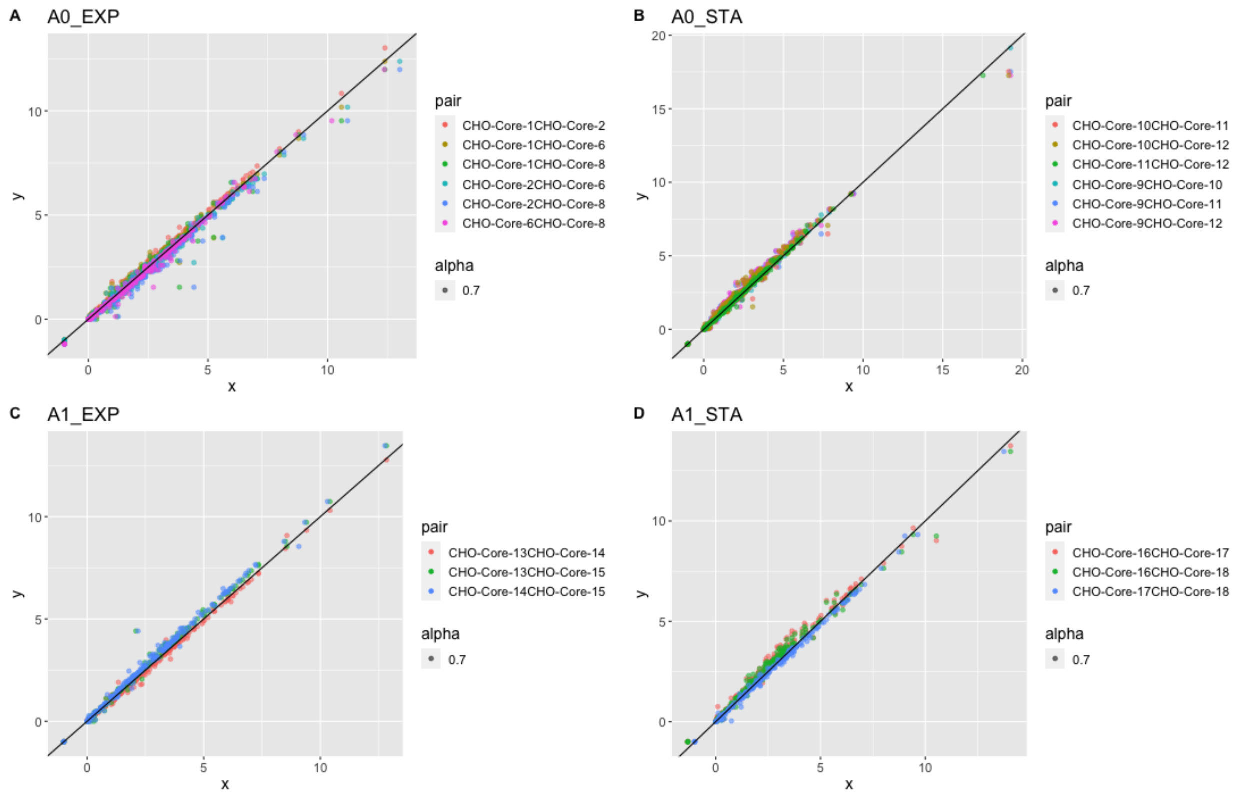


**Supplementary Figure S3.** CellFie scores of IgG producing CHO cell lines after normalization. You can see that CHO-Core-8 and CHO-Core-16 fall much closer to the reference line.

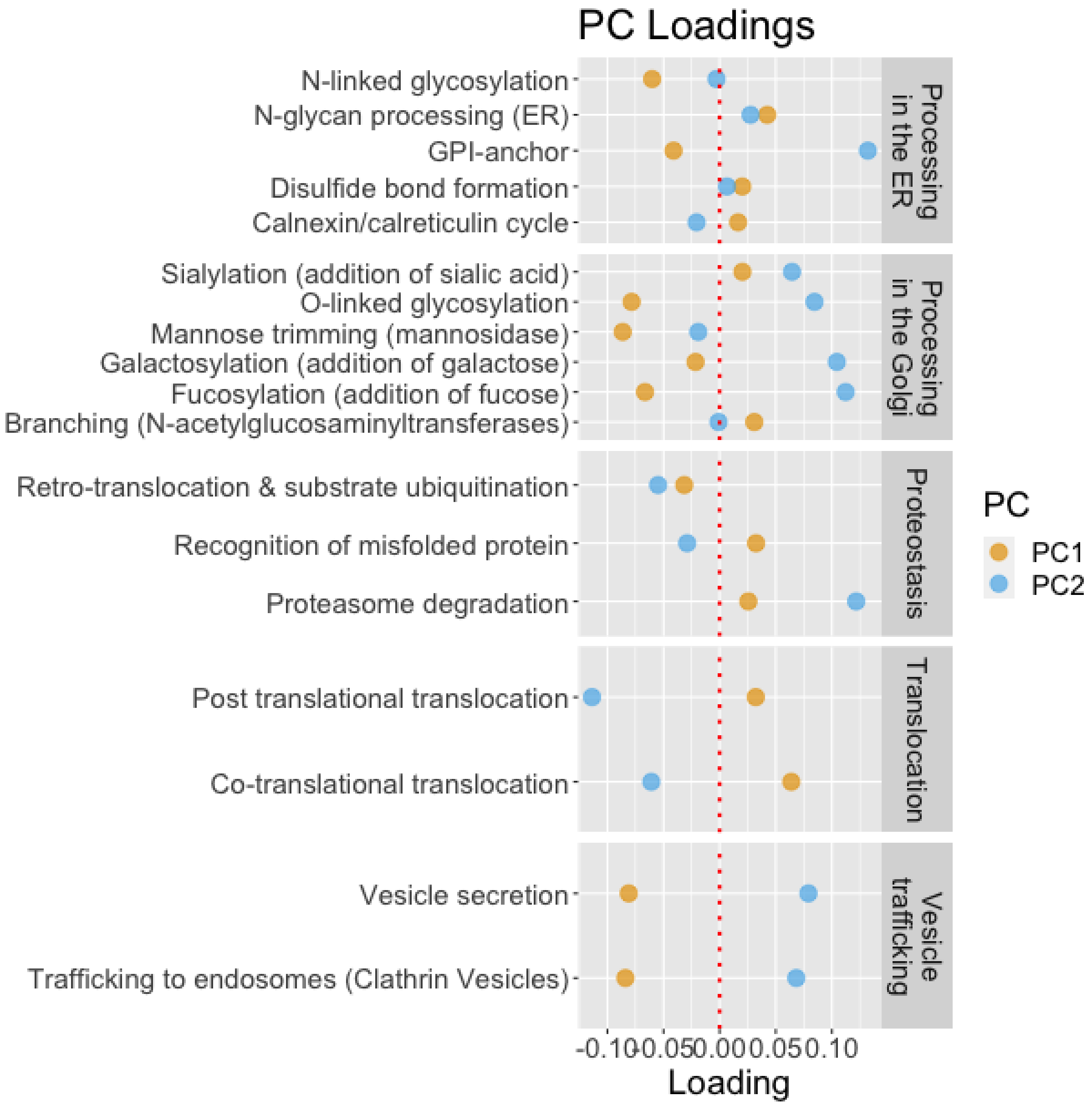


**Supplementary Figure S4.** Principal component analysis of IgG-producing CHO cells. Cleveland plot showing the effect of secretory pathway tasks on the loading of PC1 and PC2.

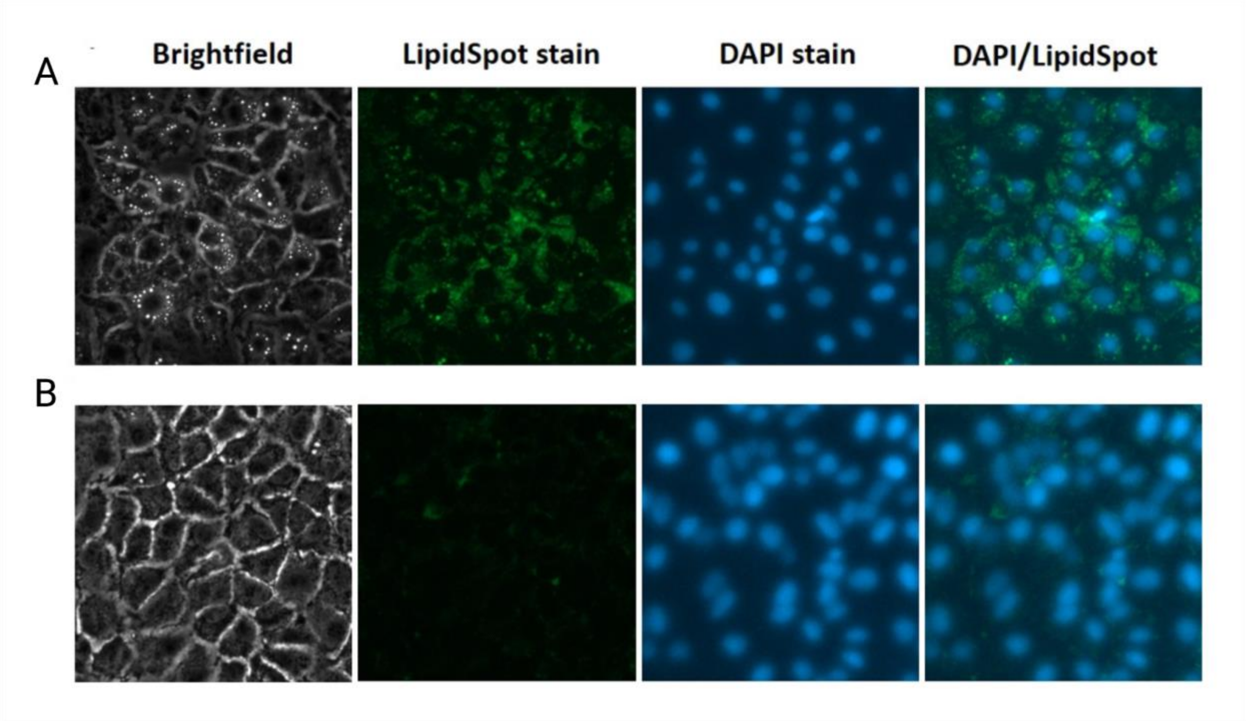


**Supplementary Figure S5. Increased lipid accumulation in NAFLD model.** Lipid droplet accumulation in Huh7 cells for **A)** fatty acid treated versus **B)** control was visualized by confocal microscopy. Wells were incubated with LipidSpot 488 for lipid staining and DAPI. Wells treated with fatty acid PA-BSA solution show increased lipid droplets by brightfield and by fluorescent staining compared to untreated cultures.

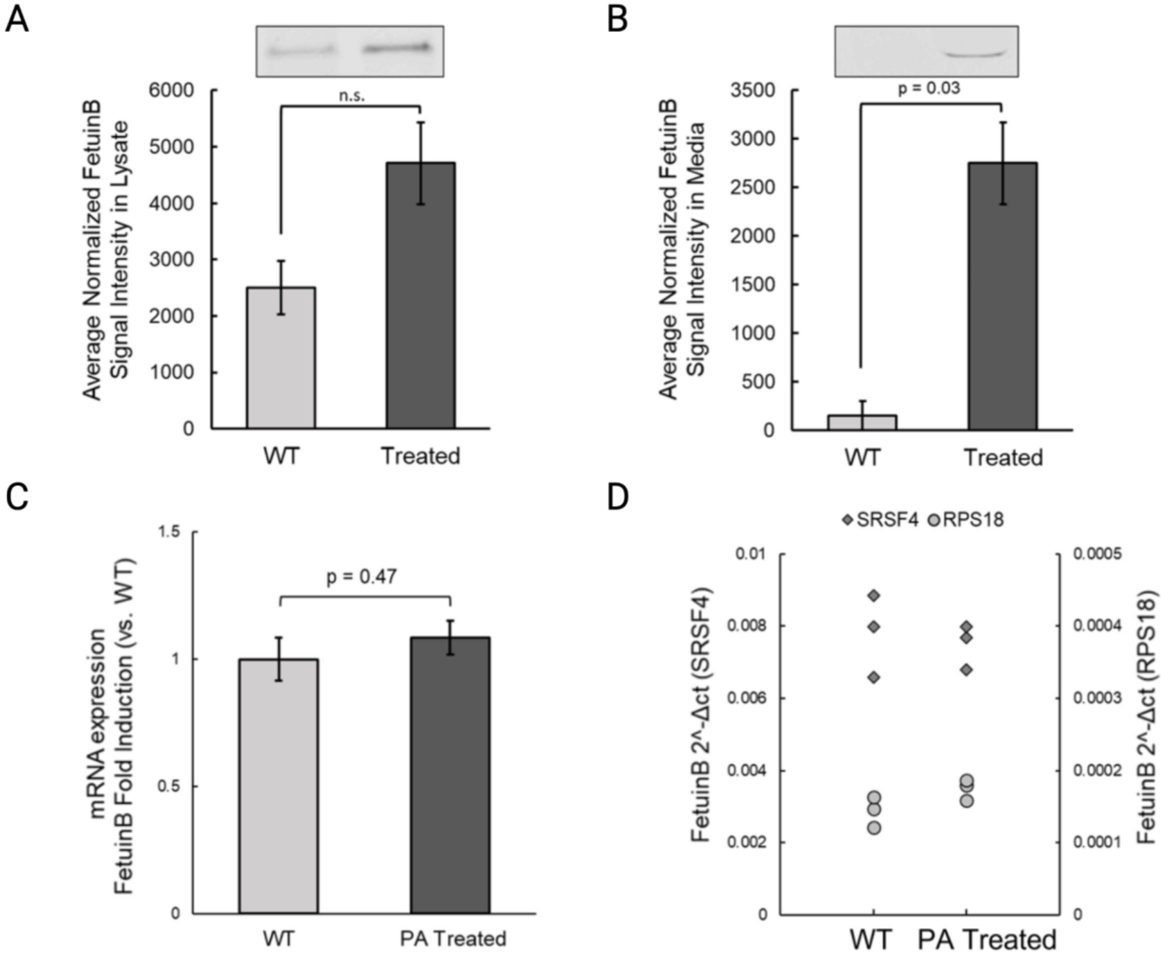


**Supplementary Figure S6. Altered fetuin-B secretion in NAFLD model.** Semi-quantitative WBs were used to measure **A)** intracellular and **B)** secreted fetuin-B from control and PA-treated culture lysate and media, respectively, from three replicates for each condition. Cultures that were fatty-acid supplimated using PA-BSA solution demonstrated increased intracellular fetuin-B and significantly increased fetuin-B within the culture media. RT-qPCR for fetuin-B using both SRSF4 and RPS18 as references was performed for triplicate samples of control and PA-treated Huh7 cell RNA extracts visualized by **C)** fold change with control and **D)** individual fold changes using controls SRSF4 and RPS18. No significant difference between transcription levels of fetuin-B between untreated and FA treated cell cultures was detected.

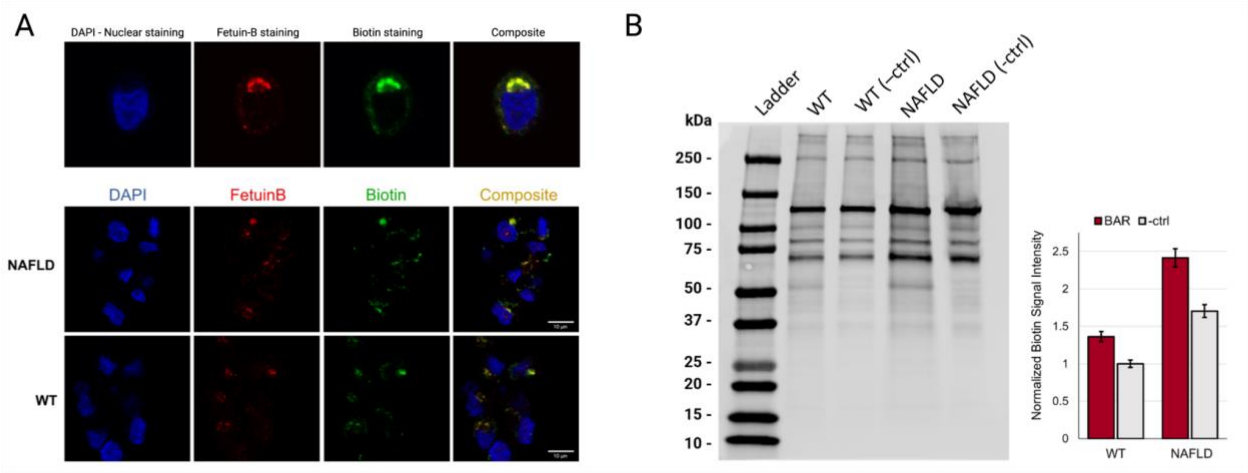


**Supplementary Figure S7. Bar effectively labels proteins proximal to fetuin-B in normal and NAFLD model cells. A)** Following biotin labeling using BAR, colocalization between fetuin-B and biotin was visualized using IF staining. **B)** WB was used to visualize biotinylation due to BAR labeling in control and NAFLD model sample lysates compared to unlabeled control (-ctrl) and NAFLD (-ctrl) Huh7 cell lysates. Performance of the BAR reaction results in biotinylation of proximal proteins to fetuin-B, which can be seen by an increase of biotinylation signal seen in the BAR treated sample lanes both for control and NAFLD. Shared bands across lanes most likely correlate to endogenous biotin-binding proteins,e.g. carboxylases that use biotin as a cofactor.

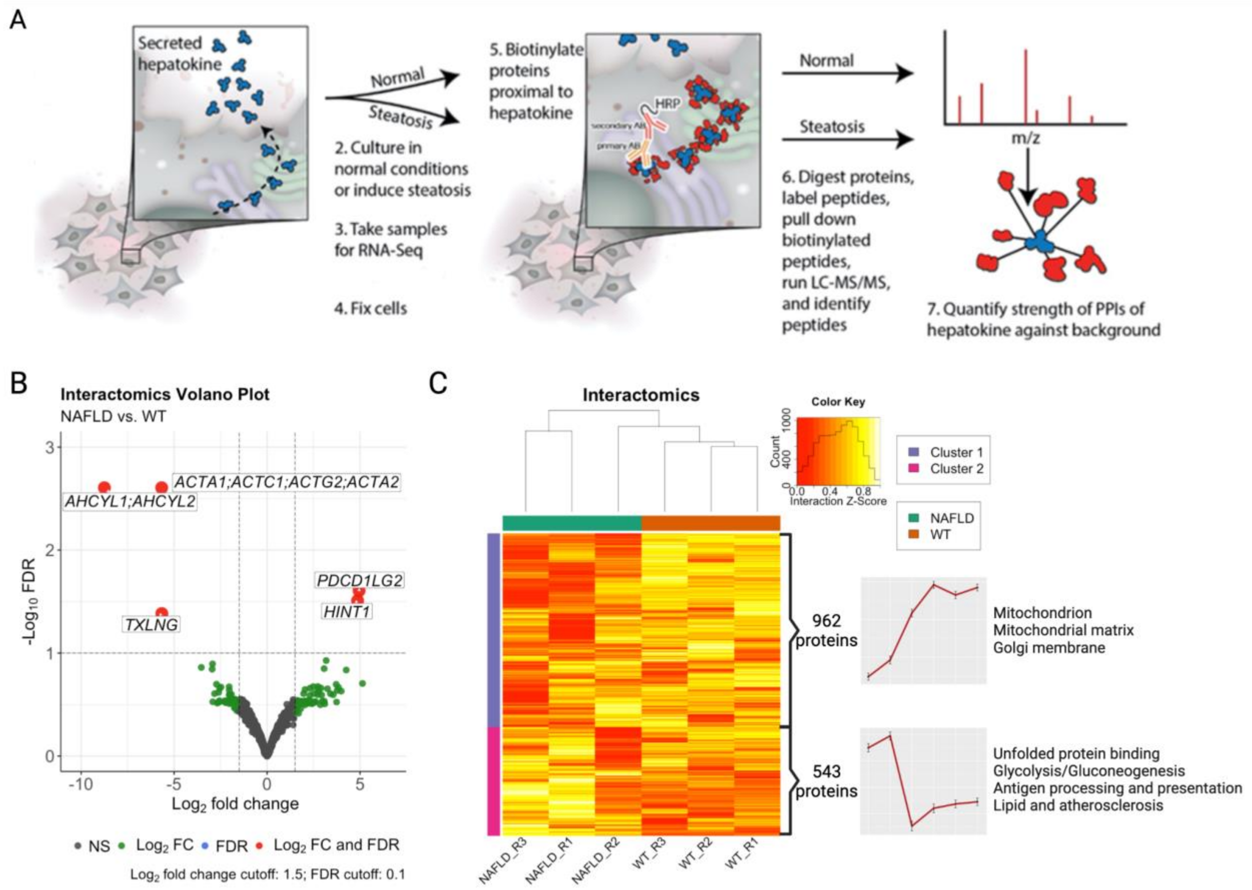


**Supplementary Figure S8. Fetuin-B interactome in NAFLD. A)** Schematic of the general experimental workflow. PPI's are measured for the model hepatokine fetuin-B using the BAR method. Relative quantification of interactions are measured in both normal and steatotic cells. **B)** Volcano plot of fetuin-B interactions measured with BAR. Interactions that passed both significance (FDR) and fold change thresholds are shown in red. **C)** Clustered heatmap of the fetuin-B interactome measured with BAR. Trend lines to the right of each cluster show mean interaction strength with the samples. Select significantly enriched terms (GO, KEGG, Reactome) for each cluster are listed on the far right.

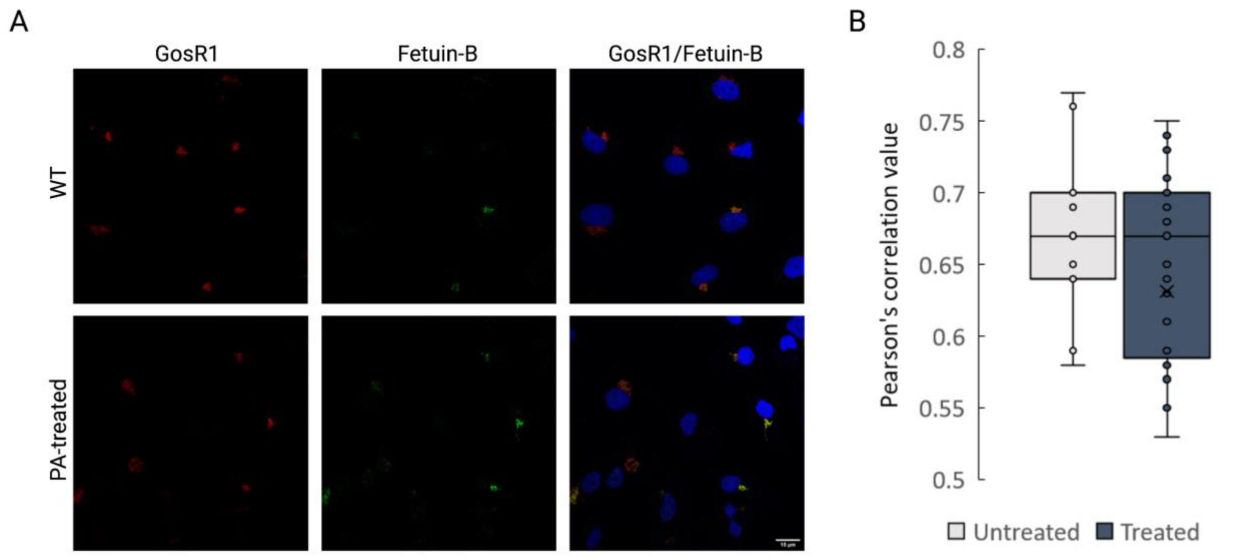


**Supplementary Figure S9. Fetuin-B co-localization with GosR1. A)** Colocalization between Fetuin-B and GosR1 visualized using IF staining for WT and PA-treated Huh7 .**B)** Pearson's correlation values of GosR1 and Fetuin-B channels for WT(n=9) and PA-treated(n=23) imaged Huh7 cells for colocalization determination. Both WT and PA-treated cells show colocalization of Fetuin-B and GosR1, suggesting that lipid accumulation does not cause Golgi-bypass during fetuin-B secretion.

Meex RC, Hoy AJ, Morris A, Brown RD, Lo JCY, Burke M, Goode RJA, Kingwell BA, Kraakman

MJ, Febbraio MA, Greve JW, Rensen SS, Molloy MP, Lancaster GI, Bruce CR, Watt MJ. 2015. Fetuin B Is a Secreted Hepatocyte Factor Linking Steatosis to Impaired Glucose Metabolism. *Cell Metab.* **22**:1078–1089.

Meex RCR, Watt MJ. 2017. Hepatokines: linking nonalcoholic fatty liver disease and insulin resistance. *Nat. Rev. Endocrinol.* **13**:509–520.

Schindelin J, Rueden CT, Hiner MC, Eliceiri KW. 2015. The ImageJ ecosystem: An open platform for biomedical image analysis. *Mol. Reprod. Dev.* **82**:518–529.
